# Clarifying space use concepts in ecology: range vs. occurrence distributions

**DOI:** 10.1101/2022.09.29.509951

**Authors:** Jesse M. Alston, Christen H. Fleming, Michael J. Noonan, Marlee A. Tucker, Inês Silva, Cody Folta, Thomas S.B. Akre, Abdullahi H. Ali, Jerrold L. Belant, Dean Beyer, Niels Blaum, Katrin Böhning-Gaese, Rogerio Cunha de Paula, Jasja Dekker, Jonathan Drescher-Lehman, Nina Farwig, Claudia Fichtel, Christina Fischer, Adam T. Ford, René Janssen, Florian Jeltsch, Peter M. Kappeler, Scott D. LaPoint, A. Catherine Markham, E. Patricia Medici, Ronaldo Gonçalves Morato, Ran Nathan, Kirk A. Olson, Bruce D. Patterson, Tyler R. Petroelje, Emiliano Esterci Ramalho, Sascha Rösner, Luiz Gustavo Oliveira Santos, Dana G. Schabo, Nuria Selva, Agnieszka Sergiel, Orr Spiegel, Wiebke Ullmann, Filip Zieba, Tomasz Zwijacz-Kozica, George Wittemyer, William F. Fagan, Thomas Müller, Justin M. Calabrese

## Abstract

Quantifying animal movements is necessary for answering a wide array of research questions in ecology and conservation biology. Consequently, ecologists have made considerable efforts to identify the best way to estimate an animal’s home range, and many methods of estimating home ranges have arisen over the past half century. Most of these methods fall into two distinct categories of estimators that have only recently been described in statistical detail: those that measure range distributions (methods such as Kernel Density Estimation that quantify the long-run behavior of a movement process that features restricted space use) and those that measure occurrence distributions (methods such as Brownian Bridge Movement Models and the Correlated Random Walk Library that quantify uncertainty in an animal movement path during a specific period of observation). In this paper, we use theory, simulations, and empirical analysis to demonstrate the importance of applying these two classes of space use estimators appropriately and distinctly. Conflating range and occurrence distributions can have serious consequences for ecological inference and conservation practice. For example, in most situations, home-range estimates quantified using occurrence estimators are too small, and this problem is exacerbated by ongoing improvements in tracking technology that enable more frequent and more accurate data on animal movements. We encourage researchers to use range estimators to estimate the area of home ranges and occurrence estimators to answer other questions in movement ecology, such as when and where an animal crosses a linear feature, visits a location of interest, or interacts with other animals.

**Open Research Statement:** Tracking data on *Aepyceros melampus, Beatragus hunteri, Bycanistes bucinator, Cerdocyon thous, Eulemur rufifrons, Glyptemys insculpta, Gyps coprotheres, Madoqua guentheri, Ovis canadensis, Propithecus verreauxi, Sus scrofa*, and *Ursus arctos* are publicly archived in the Dryad repository (Noonan et al. 2018; https://doi.org/10.5061/dryad.v5051j2), as are data from *Procapra gutturosa* (Fleming et al. 2014a; https://doi.org/10.5061/dryad.45157). Data on *Panthera onca* were taken from (Morato et al. 2018). Additional data are publicly archived in the Movebank repository under the following identifiers: *Canis latrans*, 8159699; *Canis lupus*, 8159399; *Chrysocyon brachyurus*, 18156143; *Felis silvestris*, 40386102; *Gyps africanus*, 2919708; *Lepus europaeus*, 25727477; *Martes pennanti*, 2964494; *Panthera leo*, 220229; *Papio cynocephalus*, 222027; *Syncerus caffer*, 1764627; *Tapirus terrestris*, 443607536; *Torgos tracheliotus*, 2919708; and *Ursus americanus*, 8170674.

## Introduction

Understanding how and why animals use the areas they inhabit is a core goal in the fields of ecology and conservation biology (Jeltsch *et al*., 2013, Nathan *et al*., 2008, Schick *et al*., 2008, Sutherland *et al*., 2013). The attributes of the areas where animals live shape their fitness, and knowledge of relationships between movement and fitness informs our understanding of how animals interact with each other and their environments, as well as our ability to implement effective conservation interventions (Allen & Singh, 2016). For these reasons, the importance of quantifying space use was recognized early in the development of ecology and led to the concepts of “home ranges” and “utilization distributions”. The conceptual definition of home ranges provided by Burt (1943) is still the most widely cited and targeted. Burt defined an animal’s home range as “…that area traversed by the individual in its normal activities of food gathering, mating, and caring for young. Occasional sallies outside the area, perhaps exploratory in nature, should not be considered as [a] part of the home range.” Two and a half decades after Burt offered this definition, Jennrich & Turner (1969) coined the term ‘utilization distribution’ as the probabilistic representation of a home range, providing a foundation for translating Burt’s conceptual idea into statistical estimators that can be applied to animal location data (Horne *et al*., 2020). Together, these ideas have served as the foundation of research on animal movement and resource use over the past half century.

Movement and resource use, however, are multifaceted aspects of animal behavior. Consequently, the home range concept has broadened substantially over time and there now exists a very large literature describing different approaches to home range estimation (Fieberg & Börger, 2012, Heit *et al*., 2021, Horne *et al*., 2020, Kie *et al*., 2010). Many of these approaches cluster around two distinct spatial probability distributions that arise from stochastic movement processes and can be estimated from animal location data. Fleming et al. (2015, 2016) referred to these as the “range” and “occurrence” distributions, and others have begun to adopt this terminology (Horne *et al*., 2020, Keith *et al*., 2019, Scharf *et al*., 2018, Schlägel *et al*., 2019, Signer & Fieberg, 2021). Specifically, the range distribution describes the long-run behavior of a movement process that features restricted space use and is consistent with Burt’s classical definition of the home range. In contrast, occurrence distributions quantify uncertainty in the movement path of an individual during a period of observation and are not directly related to Burt’s definition of the home range. Both of these distributions can serve as an estimation target for which specific statistical estimators can be derived, but range estimators quantify fundamentally different phenomena than occurrence estimators: range distributions answer the question “How much space does an animal need over the long term?”, while occurrence distributions answer the question “Where did an animal travel during a defined period of observation?”. Although these questions may appear similar, range and occurrence distributions have very different biological and mathematical interpretations.

In this paper, we argue that range and occurrence distributions can serve as focal points around which to organize concepts, models, statistical estimators, and research questions. We use theoretical arguments, simulations, and empirical examples to demonstrate similarities and differences between these distributions, as well as consequences that can arise from conflating range and occurrence estimators. We then link these two distributions to the ecological questions each can answer, and to the estimators that arise from each distribution.

## Concepts and Definitions

By explicitly separating the discrete-time and often arbitrary sampling schedule from the underlying continuous-time movement process, continuous-time movement models offer a number of advantages over the more traditional approach of assuming a discrete-time movement process (Kareiva & Shigesada, 1983, Langrock *et al*., 2012, Morales *et al*., 2004). These advantages include the ability to estimate scale-invariant parameters, the ability to model movement using irregularly sampled data, and freedom from the assumption of serial independence among data points (Fleming *et al*., 2014b, Gurarie *et al*., 2017, Johnson *et al*., 2008). Defining movement in this way provides a framework that facilitates the derivation of rigorous statistical procedures for quantifying movement (Blackwell, 1997, Dunn & Gipson, 1977, Fleming *et al*., 2015a, Hanks *et al*., 2015, Johnson *et al*., 2008), including many non-random behaviors such as migration, territoriality, patrolling, trap-lining, collective movement, and habitat- or condition-specific movement (e.g., Brennan *et al*., 2018, Moriarty *et al*., 2017, Papageorgiou & Farine, 2020, Péron *et al*., 2017, Sawyer *et al*., 2019). In this framework, we may consider an animal’s trajectory collected from a telemetry movement track, **r**(*t*) = (*x* (*t*), *y* (*t*)), to be a realization from a continuous-time stochastic process that is observed at discrete times *t*_1_, *t*_2_, *t*_3_, …, *t*_*n*_. From this realization, we estimate quantities related to the animal’s movement patterns, conditional upon stochastic movement models that can be used to generate movement trajectories (Table 1). Movement models such as Brownian motion (Einstein, 1905, Horne *et al*., 2007) and the integrated Ornstein-Uhlenbeck (IOU) process (Gurarie *et al*., 2017, Gurarie & Ovaskainen, 2011, Johnson *et al*., 2008) are endlessly diffusing processes and thus do not have finite coverage areas in the long run. In contrast, models such as the Ornstein-Uhlenbeck (OU; Dunn & Gipson, 1977, Uhlenbeck & Ornstein, 1930) and Ornstein-Uhlenbeck Foraging processes (OUF; Fleming *et al*., 2014a, 2015b) feature finite coverage areas, even as *t* approaches infinity. The OU and OUF processes can be thought of as range-resident versions of Brownian motion and IOU processes, respectively. Another key distinction among movement models arises from the types of autocorrelation they can accommodate. Brownian motion and OU movement produce autocorrelated positions but uncorrelated velocities, while IOU and OUF movement produce both autocorrelated positions and autocorrelated velocities. In contrast, the independent and identically distributed (IID) process, while having a finite coverage area, produces—as the name implies—completely uncorrelated data. With these movement models in mind, we can define two key families of distributions that capture many (but not all) conceptions of “space use” in the ecological literature.

**Table 1:** Summary of stochastic processes that can currently be used to model animal movement. These processes can feature positional autocorrelation, velocity autocorrelation, and/or range residency. The independent and identically distributed (IID) process can describe animal location data in which no autocorrelation is present. Brownian motion (BM) occurs in the limit of the Ornstein-Uhlenbeck (OU) process, when its positional autocorrelation time scale approaches infinity, while the Integrated Ornstein-Uhlenbeck (IOU) process occurs when the positional autocorrelation time scale of the Ornstein-Uhlenbeck Foraging (OUF) process approaches infinity. More detailed mathematical descriptions of these models can be found in Fleming et al. 2014a and Fleming et al. 2015b.

| Movement Model | Position Autocorrelation | Velocity Autocorrelation | Range Residency |
| --- | --- | --- | --- |
| IID | No | No | Yes |
| BM | Yes | No | No |
| OU | Yes | No | Yes |
| IOU | Yes | Yes | No |
| OUF | Yes | Yes | Yes |

### The Range Distribution

Movement processes that feature finite coverage areas, including the IID, OU, and OUF processes, admit a marginal distribution *p* (**r**, *t*) at each time *t*, which is the probability density of a random location **r**(*t*) being **r** at time *t*, without conditioning on any previous locations. In the most general sense, a range distribution is a marginal distribution focused on a particular time frame or suite of movement behaviors, by marginalizing over times or behaviors, to enable predictions of an animal’s locations in future periods. In other words, a range distribution describes the probability of an animal being in a location at a given time, taking into account all of the locations in a movement track simultaneously. The range distribution is simplest to define for stationary processes, which describe unchanging movement behaviors:

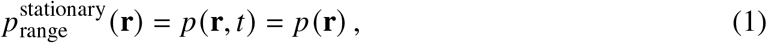

for any time *t*. Non-stationary processes, which describe movement behaviors that change over time (e.g., migrations, drifting home ranges), further require an appropriate time average to weight the relevant marginal distributions (e.g., Fleming *et al*., 2018, S1). Because 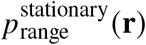 denotes the relative frequencies of different locations, the range distribution provides a prediction of *space use*, in that 95% of future locations will fall within its 95% coverage area, so long as the underlying movement process does not change (a testable assumption; see Noonan *et al*. 2019).

Range distributions therefore capture the long-run (asymptotic) area of the movement process. They are generated by running a single realization of the movement process forward into the future while keeping movement behavior fixed. The coverage areas of the range distribution are not estimates of what space the animal has used during the observation period, but predictions of what space will eventually be used, given a sufficient amount of time for the movement process to continue. All else being equal, an IID process will very quickly fill out the ranging area, whereas highly autocorrelated processes such as OUF will take longer to fill out the ranging area. However, the autocorrelation in the resulting data contains information about the long-run area of the process, and thus the *estimate* of the range distribution that accounts for autocorrelation in the data may contain a considerable amount of space that is not visited during a period of study. The range distribution corresponds closely to Burt’s conceptual definition of home range because it captures the area that the animal typically uses, not including exploratory forays. The range distribution is thus the appropriate tool for answering the question of “How large is an animal’s home range?”.

When data are statistically independent, and thus consistent with the IID assumption, the range distribution can be estimated by a variety of methods including Minimum Convex Polygons (MCPs), conventional Kernel Density Estimation (KDE), and classical Mechanistic Home Range Analysis. For the autocorrelated data provided by modern technologies such as GPS and ATLAS (Kays *et al*., 2015, Nathan *et al*., 2022), the range distribution is most accurately estimated by Autocorrelated Gaussian Density Estimation (Dunn & Gipson, 1977, Fleming *et al*., 2014b) if the home range is Gaussian, or Autocorrelated Kernel Density Estimation (AKDE; Fleming *et al*., 2015a, Noonan *et al*., 2019) otherwise. In other words, the estimation target of all of these estimators is the range distribution, but each estimator differs in the assumptions made about the data that underlie it. A given estimator must therefore be used only when the data are consistent with the movement model that underlies that estimator’s assumptions (as is standard statistical practice).

For a range distribution to exhibit a finite coverage area, the stochastic process from which it is derived must also feature finite coverage. Finite area manifests as an asymptote in the stochastic processes’ semi-variance function as the time lag between observations of the process increases (Fleming *et al*., 2014a). Some, but not all, stochastic movement models feature finite space (Table 1). These include the IID process, the OU process, and the OUF process. Importantly, as mentioned earlier, widely used models such as Brownian and IOU motion in the continuous-time context, and (correlated) random walks in discrete time, are endlessly diffusing processes and thus do not have finite range areas (Fleming *et al*., 2016). This means that these models do not provide useful estimates of home range areas.

Finally, we note that there is no dependence in the definition of the range distribution on the particular sampling regime chosen by an investigator. The range distribution is a property of the movement process that is independent of the sampling process. However, the *estimators* of the range distribution are subject to a number of biases, some of which can be related to the sampling process (Silva *et al*., 2022). First, a range estimate becomes more fully resolved in proportion to its “effective sample size”, which is approximately how many times the focal animal crossed its home range during the observation period. If the animal has not crossed its range during the observation period, it is not possible to estimate the range distribution. Second, different estimators of the range distribution may exhibit either positive or negative biases that decrease asymptotically as sampling duration increases. Third, estimators that assume IID data (e.g., conventional KDE, MCP, Mechanistic Home Range Analysis) tend to underestimate the ranging area when applied to autocorrelated tracking data by an extent that depends, all else equal, on the strength of autocorrelation in the sampled locations (Noonan *et al*., 2019). Again, this is not an inherent property of range distributions *per se*, but, instead, results from using estimators for which a core assumption has been violated. As with any statistical procedure, violating a key assumption of a home range estimator can produce biased results.

### The Occurrence Distribution

Whereas range distributions are based on the marginal distributions *p* (**r**, *t*) and can predict unrealized locations, occurrence distributions are based on the conditional distributions *p* (**r**, *t*|data) and are focused on interpolating movement tracks between known locations during an observation period. In other words, an occurrence distribution describes the probability of an animal being in a location at a given time, conditional upon its previous and subsequent locations. Such conditional distributions exist for all stochastic movement processes, even when those processes do not have finite coverage areas in the long run and do not describe range-resident movement behaviors (e.g., Brownian motion and IOU movement). The simplest occurrence distribution that we can construct involves uniformly averaging these conditional distributions over the observation period for times sampled between *t*_1_ and *t*_*n*_:

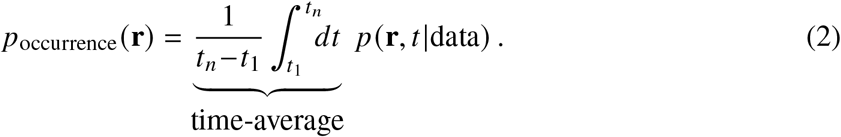

This corresponds to the conditional distribution of a realized location **r**(*t*) at a random time *t* within the observation window. However, missing observations are often skipped to avoid oversmoothing (e.g., Bedrosian *et al*., 2018, Coe *et al*., 2015, Sawyer *et al*., 2009), and one could envision a more rigorous weighting scheme that maintains a balance between detail and continuity. In the limit of very coarse, uncorrelated data, and with some gap-skipping heuristic applied, the occurrence distribution reduces to the empirical distribution of the data. This means that *there must be autocorrelation between data points* for an occurrence estimator to perform well (i.e., to narrow down the area an animal may have traveled between known locations). Estimating an occurrence distribution using data that is so coarse as to be IID, or nearly so, will provide little information on the movement track of an animal.

The occurrence distribution quantifies where an animal may have traveled during the observation period given the observed data, and relies on an autocorrelated movement model to interpolate the data. The occurrence distribution’s area is generated by considering all possible trajectories that are consistent with the data, weighted by their probability density. As the movement path of an animal becomes more finely and more accurately resolved, this area becomes smaller and smaller, eventually limiting to zero, even though actual space used has not changed. The area of occurrence estimates therefore does not directly measure space use—even during the observation period— but is, instead, a reflection of our uncertainty regarding where an animal was located during an observation period. In other words, if we have complete knowledge of the animal’s locations during an observation period (i.e., infinite sampling rate and no location error), the occurrence distribution collapses to the animal’s movement path and has zero area. The occurrence distribution is thus appropriate for answering questions such as “Where might an animal have traveled during an observation period?” and “What landscape features might an animal have visited along its movement path?”.

The occurrence distribution is not well-estimated by the range estimators outlined in the prior subsection, and proper occurrence estimators have not been around nearly as long as range estimators—occurrence estimators were introduced in the peer-reviewed ecology literature only around 15 years ago (Horne *et al*., 2007). Currently, Brownian bridge movement models (BBMMs; Horne *et al*., 2007, Kranstauber *et al*., 2012), the Correlated Random Walk Library (CRAWL; Johnson *et al*., 2008), and the generalized time-series Kriging framework (Fleming *et al*., 2016) all share occurrence distributions as estimation targets. Note that the Kriging framework contains both the BBMM and CRAWL as special cases—Kriging with a Brownian motion model is equivalent to the BBMM, while Kriging with an IOU process is equivalent to the model used in CRAWL (Fleming *et al*., 2016). The occurrence distribution exists for any autocorrelated movement process, whether or not the focal process features finite coverage areas. This means that the Brownian motion, IOU, OU, and OUF continuous-time processes all admit occurrence distributions. For an IID process, the occurrence distribution is simply the empirical distribution with some heuristic to account for gaps in the data.

Transitioning from marginal distributions that are independent of specific events to conditional distributions that are conditional upon preceding and subsequent events has a dramatic effect on the meaning and operation of occurrence distributions. Range distributions and their constituent marginal distributions are parameters of the movement process that exist independent from the sampling process (though *estimators* of the range distribution may exhibit some sampling dependence). In contrast, occurrence distributions are conditional upon the observed data and are thus explicitly defined in terms of the sampling schedule. This means that a different sampling schedule applied to the same movement process will *correctly* yield a different occurrence distribution: all else equal, increasing the sampling rate will result in a narrower, more concentrated occurrence distribution. This happens because more frequent sampling more fully resolves the animal’s true movement path, and thus uncertainty in the animal’s locations decreases concomitantly. It is important to realize that this is not due to sampling-dependent bias of occurrence estimators: occurrence estimators in the time-series Kriging family, including the BBMM, can be unbiased. Instead, the uncertainty decreases because the estimation target itself (i.e., the occurrence distribution) is a function of the sampling schedule. Figure 1 shows this process occurring for data from a fisher (*Pekania pennanti*) tracked for 19 days in New York, USA, at a roughly 2-minute sampling interval.

**Figure 1:**
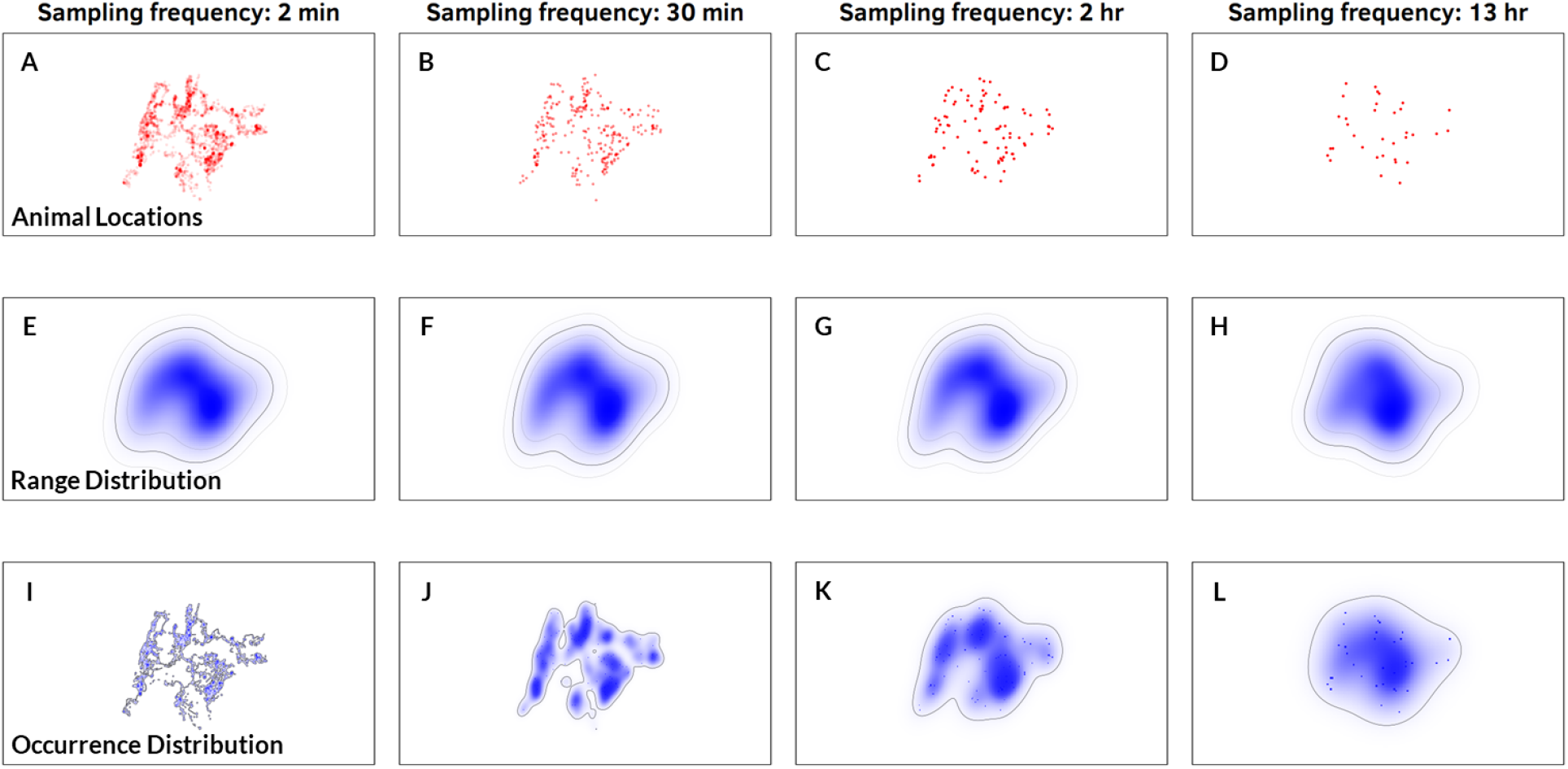
Demonstration of sampling dependence of occurrence and range distributions using GPS location data from a GPS-tracked fisher (Pekania pennanti) from New York, USA. The fisher was tracked for 19 days at 2-minute intervals. The top row features individual locations along the fisher’s movement track as the movement track is progressively thinned from 720 locations per day to 2 locations per day. The second row features 95% AKDE range estimates generated using the same GPS locations. While the contours of the range estimate change as the data are more finely resolved, the area within those contours remains largely stable. The third row features 95% Kriged occurrence estimates generated using the same data. In contrast with range estimates, the area of occurrence estimates shrinks rapidly as the data are sampled more frequently and the fisher’s movement path is more accurately resolved.

### Relationships Between Range and Occurrence Distributions

As detailed above, the range and occurrence distributions are based on different biological and statistical definitions, have different interpretations and statistical estimators, and respond differently to variation in sampling schedules. We now consider two key limits defined by data amount and quality that highlight the conditions under which range and occurrence distributions either converge or diverge completely, and reiterate a conceptual difference between the two distributions.

#### Convergent Limit: Infinite Observation Period

Given an infinite observation period, the occurrence distribution will limit to a distribution close to the range distribution, but with an amount of estimation error determined by location error and the sizes of gaps in the data. This happens because an animal visits more and more of its home range over time. Decreasing location error and increasing the sampling rate will reduce this estimation error, but increasing the sampling rate will also slow down convergence, because the occurrence area limits to zero if the sampling rate is infinite while the observation period is finite.

#### Divergent Limit: Infinite Sampling Rate

For the occurrence distribution of any real movement process that is continuous in both location and velocity, holding the sampling duration constant while increasing the sampling rate with either no location error or uncorrelated location error yields the limit:

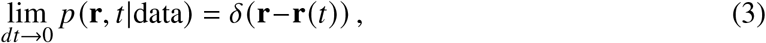

where *δ*(**r**) is the Dirac delta function—a singular distribution with probability mass concentrated at **r**. This limit is easiest to see in the case of a Brownian bridge, where the width of the bridge is at most proportional to *dt*. In any case, the occurrence distribution collapses toward the movement path even in the presence of (uncorrelated) location error as sampling becomes finer and finer, eventually collapsing to zero area. The range distribution is unaffected by this limit and its area remains the same, though estimators of the range distribution may exhibit varying sampling dependence. Increasing the rate of sampling results in increasingly strong autocorrelation in the data, so asymptotically consistent range estimators that do not account for this autocorrelation perform worse as sampling rate increases. Such estimators are increasingly negatively biased by increasing autocorrelation strength and will also limit to zero area. However, range estimators that properly model autocorrelation will be unaffected by this limit, and their area estimates will remain consistent.

#### Interpolation vs. Extrapolation

Another way of distinguishing between range and occurrence distributions is in terms of the statistical operations to which they conform. Given a sample of tracking data of finite duration, the range distribution represents an *extrapolation* of the long-run behavior of the movement process, as inferred from the data, and quantifies the variance of the movement process. In contrast, the occurrence distribution *interpolates* within the observation period, conditional on the data and an autocorrelated movement model, and quantifies uncertainty in the interpolation. This is why the general framework for occurrence estimation is based on Kriging, which is a statistically optimal method of model-based interpolation (Fleming *et al*., 2016).

To illustrate this more concretely, consider cross-validation of home range estimators. If an estimator accurately quantifies an individual’s home range (*sensu* Burt, 1943), an unbiased 95% home range area estimate generated over some observation period *T*_1_ should contain, on average, 95% of that animal’s locations over a subsequent observation period *T*_2_, provided the animal’s movement behavior does not meaningfully change between the training (*T*_1_) and test (*T*_2_) sets, and provided that *T*_1_ and *T*_2_ begin far enough apart to be uncorrelated. If a 95% home range area estimate were to consistently include more than 95% of the subsequent locations, then estimates would be positively biased; if estimates consistently include fewer than 95% of the subsequent locations, then estimates would be negatively biased. Similarly, a 50% home range estimate generated over *T*_1_ should contain 50% of an animal’s locations, on average, over *T*_2_. In other words, the relevant test set for a range estimator is an animal’s movements *in the future*, extrapolated from past location data. This can be achieved via half-sample cross-validation (e.g., Noonan *et al*., 2019). Cross-validation of occurrence estimators should operate differently. If an estimator captures an occurrence distribution accurately, an unbiased 95% area estimate generated over *T*_1_ should contain 95% of that animal’s locations within *T*_1_. In other words, the relevant test set for an occurrence estimator are holdout data from within the initial observation period (and not a subsequent period).

## Simulated Examples

The two limits described above are crucial for understanding the differences between occurrence and range distributions. We now demonstrate the importance of these limits with both simulated and real data. For the simulated data, we can specifically model processes where both types of distributions exist: processes that are (1) autocorrelated (so that the occurrence distribution can interpolate the data) and (2) range-resident (so that the range distribution exists). Simulation also allows us to set the true size of the home range (range distribution). We can then manipulate the sampling schedule of the simulated processes to explore the effects of sampling rate and sampling duration on the sizes of range and occurrence estimates. To do this, we simulated movement paths from an OUF process while varying the sampling rate and sampling duration systematically to illustrate differences between estimates provided by range and occurrence estimators. For each data set, we estimated the range distribution via Autocorrelated Gaussian Density Estimation conditioned on a fitted OUF model with daily autocorrelation timescales. Similarly, we estimated the occurrence distribution for each data set by Kriging with an OUF model with daily autocorrelation timescales. The assumptions of these estimation approaches exactly match the process that generated the data, so the statistical estimators are correctly specified for both the range and occurrence distributions in these simulated examples.

Figure 2A shows the area of the occurrence estimate decreasing as the sampling rate increases from one observation per day to 128 observations per day. Note that the 95% occurrence area starts substantially less than the range area, because the sampling duration is finite (256 days, in this case), and then rapidly collapses to zero as the sampling rate increases. Ongoing technological advances that facilitate ever finer and more accurate location sampling are driving movement studies closer to the limit where estimates produced by occurrence estimators collapse to zero area. It is therefore inevitable that the differences between range and occurrence distributions will become more obvious in the future, even though these distributions have been frequently conflated in the past.

**Figure 2:**
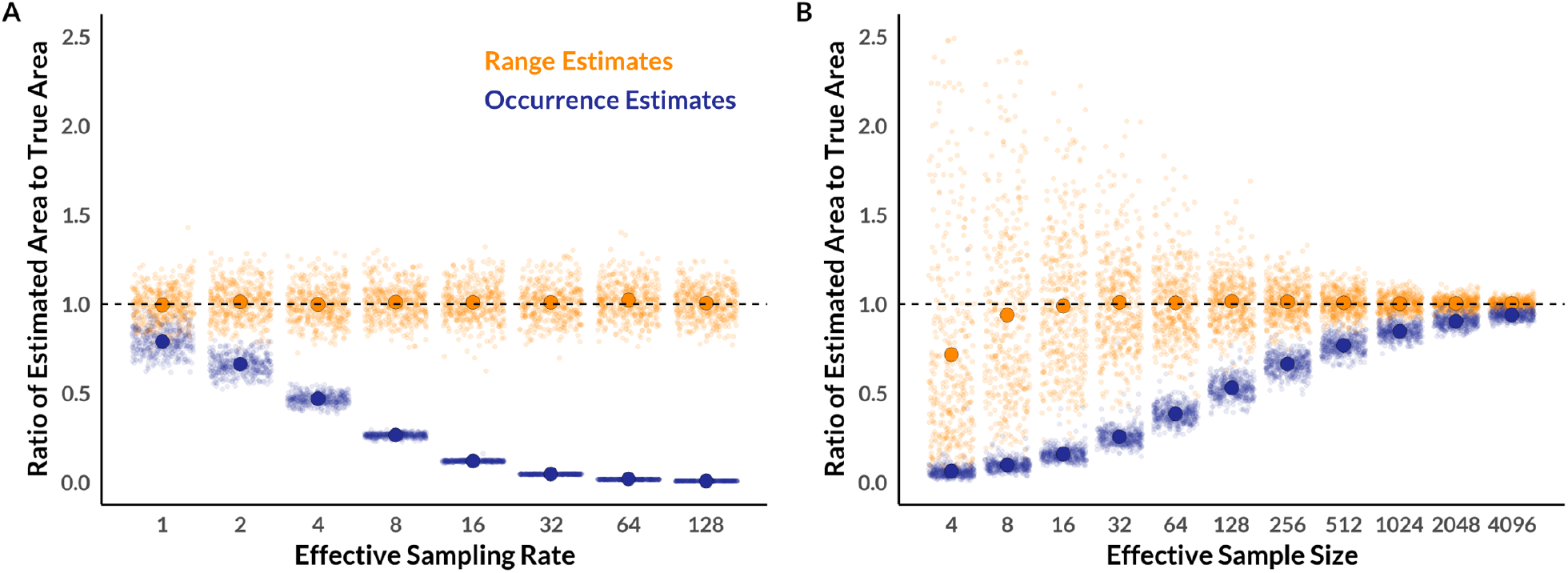
Bias in estimates of home range size provided by range (orange) and occurrence (indigo) distribution estimates with different sampling rates (i.e., GPS fixes per range crossing; Panel A) and effective sample sizes (i.e., number of range crossings in a data set; Panel B). A point at 1 indicates that home range size was estimated correctly for a simulated movement track with a known home range size. Small points represent a single simulation result (jittered on the x-axis to ease visualization), while larger points represent the mean simulation result among 400 replicates. Divergence of range estimates from the line at 1 at small durations of sampling arise from known patterns of bias that can be improved by bootstrapping (Fleming et al., 2019), while bootstrapping does little to change the size of occurrence estimates.

Figure 2B shows the area of the occurrence estimate increasing as the sampling duration increases from 4 days to 4,096 days. Again, the 95% occurrence area is still substantially less than the range area even when the effective sample size (i.e., the number of range crossings) is > 4,000, and rapidly collapses to zero as the duration of the observation period decreases. This is a major real-world problem because it can take weeks or months on average for an animal to cross its range (depending on species), and the lifespan of tracking devices (or even animals) is unlikely to enable an effective sample size of anywhere near 4,000 (i.e., 11 years with daily range crossings). This demonstrates that while occurrence distributions tend toward range distributions as the sampling duration increases, using an occurrence estimator to quantify the size of home ranges will yield a substantial underestimate unless very large amounts of data are collected—amounts that are likely logistically and/or biologically impossible. Although technological advances are increasing the battery lifespan of animal tracking devices, and thus the potential duration of animal tracking studies, the duration of tracking data for an individual animal is often limited in practice by mortality or equipment failure. Occurrence estimators will therefore tend to provide home-range estimates that are substantially smaller than the true home range size in most real-world situations.

## Empirical Examples

Using empirical data, we now show how profoundly range and occurrence estimates can diverge in real-world data sets. As outlined above, this happens when the data are sampled frequently enough that the occurrence distribution collapses toward the movement path and for long enough that estimating the range distribution is possible. Such data sets are already common and their availability will only increase as tracking technology improves (Gupte *et al*., 2022, Kays *et al*., 2015, Nathan *et al*., 2022). Using a data set of 369 individual animals across 27 species (Noonan *et al*., 2019), we estimated both the range and occurrence distributions for each animal. We estimated the range distribution via Autocorrelated Kernel Density Estimation (AKDE) conditioned on a fitted movement model according to the workflow described in Silva *et al*. (2022). In short, we used variogram analysis (Fleming *et al*., 2014a) to ensure animals were range-resident, fit and selected an autocorrelated movement model that best described the animal’s movements using perturbative Hybrid Residual Maximum Likelihood (phREML; Fleming *et al*., 2019) and Akaike’s Information Criterion corrected for small sample sizes (AICc), and estimated weighted AKDE utilization distributions for each animal (Fleming *et al*., 2018) using the `ctmm` R package (v0.6.2; Calabrese *et al*., 2016) in the R statistical software environment (v3.6.2; R Core Team, 2020). We estimated the occurrence distribution for each animal based on Kriging (Fleming *et al*., 2016) with the same movement model used for the corresponding AKDE estimate. Figure 3 shows that the occurrence estimate is smaller than the range estimate for the vast majority of individuals in the data set (and usually much smaller). This occurs because the effective sample size is rarely large enough in real-world data to approach the theoretical limit where the occurrence distribution would converge with the range distribution.

**Figure 3:**
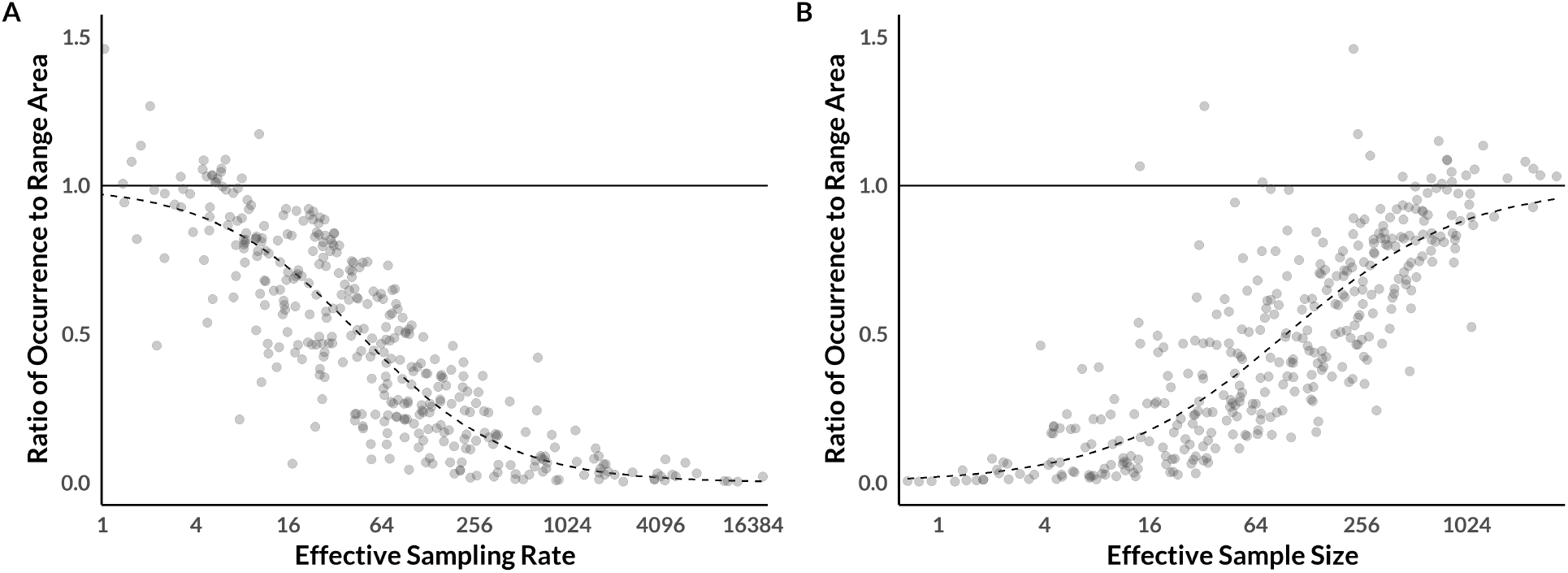
The ratio of the size of the occurrence estimate to the size of the range estimate for a data set containing 369 individuals across 27 species, as a function of the effective sampling rate (i.e., GPS fixes per range crossing; Panel A) and effective sample size (i.e., number of range crossings in a data set; Panel B). Points represent individual animals, while dashed lines represent regressions demonstrating the overall trend. Solid horizontal lines indicate a ratio of 1:1, where range and occurrence estimates are the same size. Distance below the solid line indicates the extent to which occurrence estimators are negatively biased in their estimates of home range size.

This shrinkage of occurrence estimates is accompanied by a decreased ability of the occurrence distribution to correctly specify the areas of home ranges. To illustrate this, we performed half-sample cross-validation on the same animal location data set. We subset data from each individual animal into halves, used the first half of the data to generate range (AKDE) and occurrence (Kriging) estimates, and then used the second half to assess the percentage of future animal locations that were within the range and occurrence estimates. All data fit the assumptions of range-resident animals with movement processes that remained consistent between the two halves. We then fit regression lines (linear for AKDE estimates, logistic for Kriging estimates) for the influence of effective sampling rate (roughly the number of GPS locations per range crossing) and effective sample size on the percentage of locations in the test set that fell within estimates generated using the training set.

As these results show (Fig. 4), the areas of home ranges derived from occurrence distributions do not merely fit the data more tightly—they inaccurately represent the true area of home ranges. Estimates of home ranges produced by occurrence estimators are nearly always too small, and this negative bias is exacerbated at high sampling rates and low effective sample sizes. This is not merely a hypothetical problem, nor is it only an issue that will arise in the future as technology continues to improve. Instead, it is pervasive in the animal movement data that wildlife biologists currently collect and analyze (Noonan *et al*., 2020, 2019).

**Figure 4:**
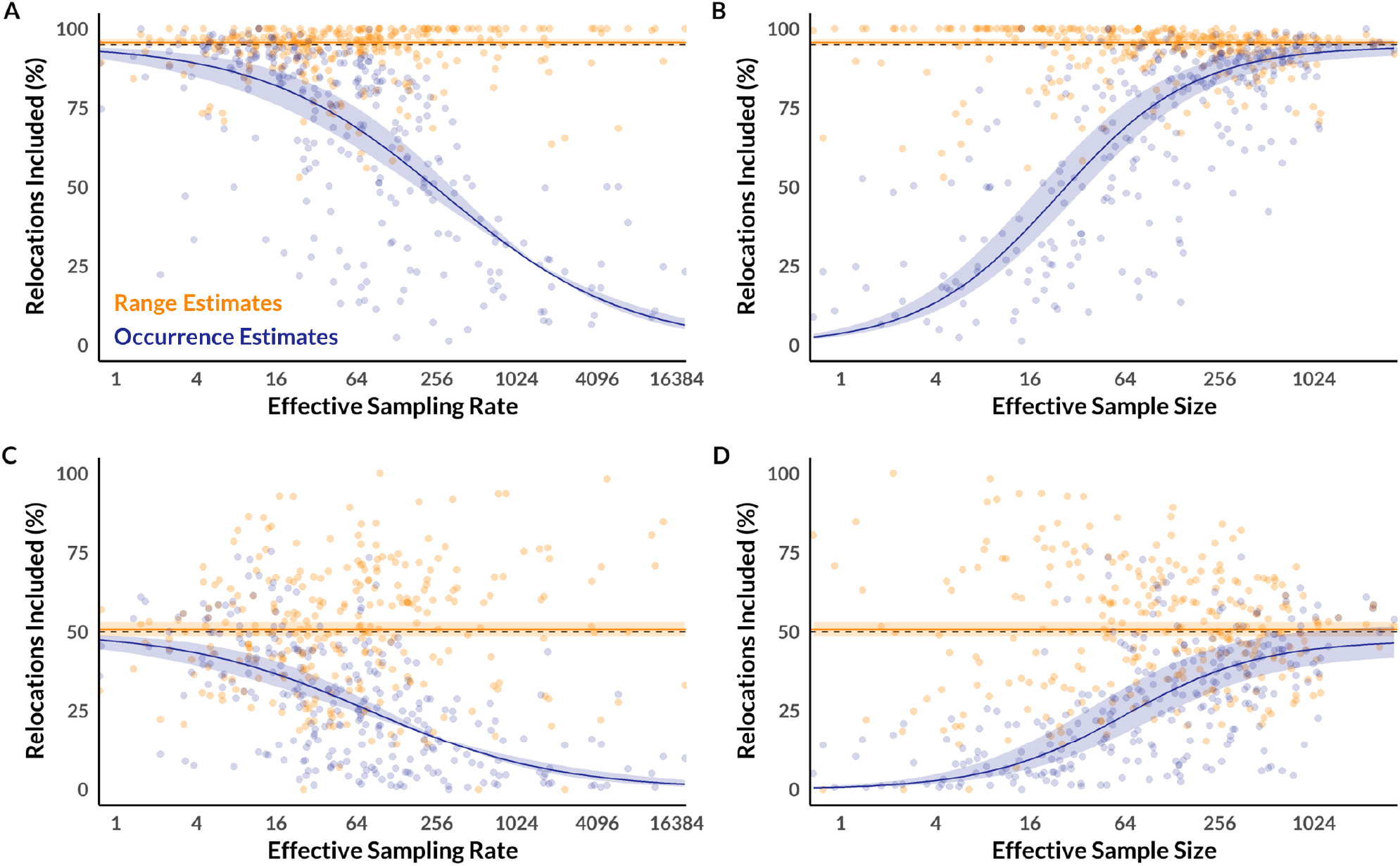
Half-sample cross-validation of range and occurrence estimates. Points represent the percentage of locations from the second half of the data (test set) included in home ranges estimated from the first half of the data (training set). Orange points represent range (AKDE) estimates, while indigo points represent occurrence (Kriging) estimates. The dashed line represents the target 95% (top row) or 50% (bottom row) quantile, while the solid line represents a regression model fit to the cross-validation results with shading to indicate the 95% confidence interval. The left column demonstrates the influence of effective sampling rate on cross-validation results, while the right column demonstrates the influence of effective sample size on cross-validation results. On average, range estimates contain roughly the correct percentage of relocations, and this remains true across all effective sampling rates and effective sample sizes. Occurrence estimates, however, tend to contain too few relocations, and this problem is exacerbated at high effective sampling rates and low effective sample sizes.

## Discussion

Ecologists often conflate occurrence estimators with range estimators, a much older and more familiar class of statistical tools. The first widely used occurrence estimators (Horne *et al*., 2007, Johnson *et al*., 2008) were landmark advances in movement ecology and enabled more statistically rigorous analyses of many research questions related to animal movement. Nevertheless, although they have been widely used in movement ecology, the extent of their novelty and unique properties have still largely gone unrecognized. As we have demonstrated, these two classes of estimators have radically different properties, and should therefore be used for different purposes (Table 2). Ecologists and conservation biologists should use range estimators to estimate the area of home ranges, and occurrence estimators to answer other questions, such as: Where might an animal have crossed a linear feature (Find’o *et al*., 2018, Hooker *et al*., 2020, Zeller *et al*., 2018)? How likely is it that an animal visited a location of interest (Noonan *et al*., 2018, Pagès *et al*., 2019, Sasmal *et al*., 2019)? When and where could two individual animals have interacted (Schlägel *et al*., 2019)? Which areas of a landscape contain high-priority resources (e.g., migratory corridors or stopover sites; Sawyer *et al*., 2009, 2019)?

**Table 2:** Summary of the primary distinctions between range and occurrence distributions.

|  | Range Distribution | Occurrence Distribution |
| --- | --- | --- |
| Distribution Type | Marginal | Conditional |
| Finite Coverage Area | Arises only when the stochastic movement process being modeled has a finite coverage area | Arises when the sampling rate is finite, regardless if the stochastic movement process being modeled has a finite coverage area |
| Statistical Operation | Extrapolation (Over what area is an animal likely to range in the future?) | Interpolation (At which locations might an animal have occurred in the past?) |
| Sampling Dependence | No (If its statistical assumptions are met, a range estimator will estimate a stable area even as a movement track is sampled more frequently) | Yes (If its statistical assumptions are met, an occurrence estimator will estimate a smaller area as a movement track is sampled more frequently) |
| Appropriate Questions | How large is an animal's home range? What area is available to an animal in studies of third-order habitat selection? | Where might an animal have crossed a linear feature? When and where could two individual animals have interacted? In which areas of a landscape did an animal visit high-priority resources (e.g., migratory corridors or stopover sites)? |

Estimation of animal home ranges is foremost among our concerns on the conflation of range and occurrence estimators—occurrence estimators substantially underestimate the area of home ranges under a broad array of real-world conditions (Figs. 3,4). In recent years, there has been a slow but steady drift in preference among wildlife biologists towards estimators that fit more tightly to animal location data (Fig. 5; Crane *et al*., 2021, Laver & Kelly, 2008, Walter *et al*., 2015). We believe that this preference has largely been driven by aesthetic considerations and the intuitive notion that areas within home range estimates where an animal does not travel during a study are not actually “used” (Cumming & Cornélis, 2012, Getz *et al*., 2007, Kie, 2013, Walter *et al*., 2015). This preference can be observed in the transition over time from home range estimates using Minimum Convex Polygons to Local Convex Hull (LoCoH; Getz *et al*., 2007, Getz & Wilmers, 2004) to Time Local Convex Hull (T-LoCoH Lyons *et al*., 2013) methods, an emphasis on KDE bandwidth optimizers that fit tightly to location data (Cohen *et al*., 2018, Downs & Horner, 2008, Kie, 2013), and most recently, rapid growth in use of BBMMs to estimate home ranges (Fig. 5). Cross-validation frameworks that seek to backtest estimator performance (e.g., Getz & Wilmers, 2004, Kie, 2013, Silva *et al*., 2020, Walter *et al*., 2015), which are appropriate for occurrence estimators but not range estimators, have also provided a false impression that smaller home range estimates perform better. While understandable, seeking home range estimates that fit tightly to an animal’s past locations adheres neither to Burt’s original definition, nor the mathematical properties underlying the range distribution. Specifically, Burt’s definition aims to capture the amount of space an animal will need to survive and reproduce in the long run, not simply the level of uncertainty in an animal’s movement path during an observation period limited by study design, technology, or animal mortality.

**Figure 5:**
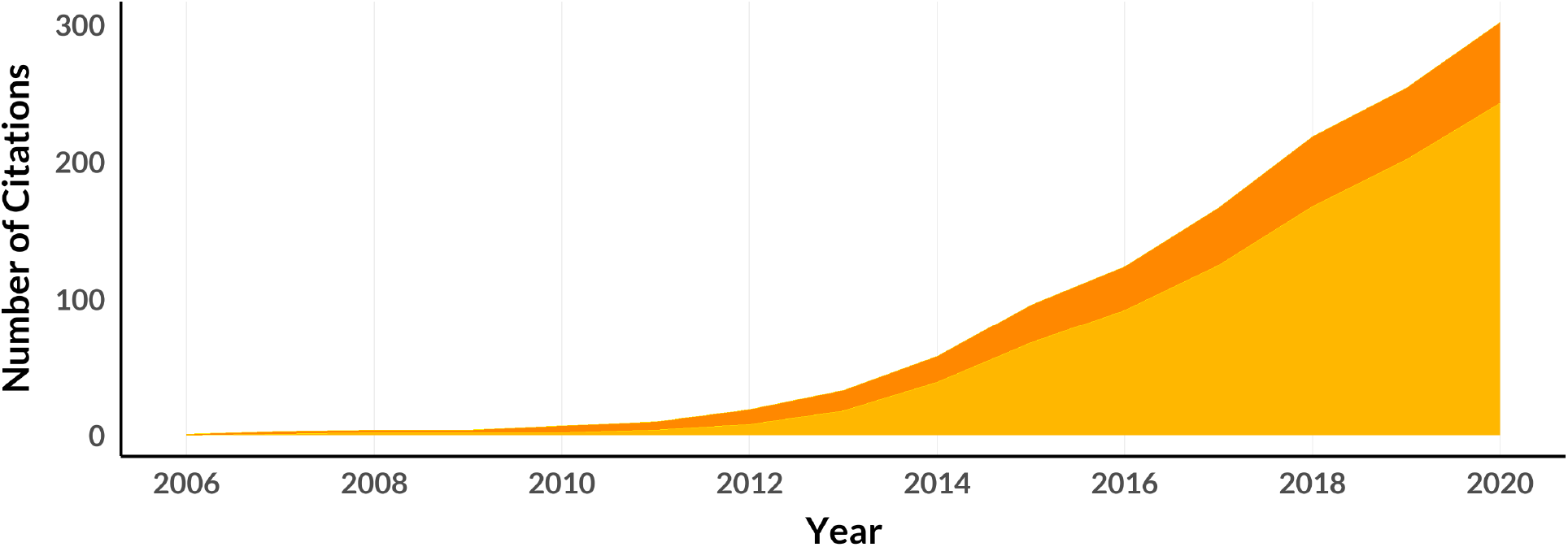
Number of peer-reviewed journal articles from 2006 to 2020 that either used BBMMs to estimate the size of animal home ranges (light orange) or labeled BBMMs as a home range estimator (dark orange). Although BBMMs are occurrence estimators and therefore poorly suited for estimating the size of home ranges, use of BBMMs to estimate the size of home ranges is growing rapidly.

The most common occurrence estimator used to estimate home ranges is the BBMM (Horne *et al*., 2007), which has even been championed as a “third generation home range estimator” due to its ability to account for some autocorrelation in tracking data (Walter *et al*., 2015). Figure 5 shows the cumulative number of peer-reviewed journal articles since the BBMM was introduced to ecologists in 2007 that either label it a home range estimator, or use it to estimate animal home range areas. However, Fleming et al. (2016) formally proved that the BBMM is an estimator of the occurrence distribution (rather than the range distribution) that arises as a special case of the more general time-series Kriging family of occurrence estimators. Specifically, Kriging a movement track conditional on a Brownian motion movement model is equivalent to the BBMM. Beyond being an occurrence estimator, and thus only suited to the task of home range estimation in the (unrealistic) infinite data limit, the BBMM is also based on an endlessly diffusing Brownian motion process, which does not have a finite range area. Note that this is not a critique of the validity of the BBMM as an analytical tool *per se*. Other occurrence estimators, such as a time-series Kriging estimate based on an OUF process, are also inappropriate for estimating the area of home ranges (as demonstrated in Figs. 1-3), and BBMMs are the best tool currently available for quantifying where an animal might have been during an observation period if the Brownian motion model accurately characterizes the animal’s movement process. Furthermore, when BBMMs were developed, the issue of underestimation of home ranges as outlined above was not as apparent, because animal location data were coarser than they are today. However, as animal tracking technology improves and the resolution of data sets increases, the discrepancy between the area that BBMMs estimate and a proper estimate of the range distribution will continue to widen and repeated studies of the same species with improved technology will lead to progressively smaller estimates of home ranges if these estimates are generated using occurrence estimators like BBMMs.

Using occurrence estimators to quantify home ranges can therefore have pernicious consequences for area-based conservation strategies and for ecological inference. For example, many protected areas (e.g., the Attwater Prairie Chicken National Wildlife Refuge, Kirtland’s Warbler Wildlife Management Area, and the National Key Deer Refuge in the USA, the Arawale National Reserve in Kenya, and the Blackbuck Conservation Area in Nepal) are designed to protect a focal species. For these protected areas, understanding how much space is required to maintain populations that are viable over the long term is vital for ensuring their effectiveness (Brashares *et al*., 2001, Pe’er *et al*., 2014). When protected areas are too small relative to their focal species’ area requirements, the probability of population declines or extirpation increases significantly (Brashares *et al*., 2001, Gaston *et al*., 2008). Undersized protected areas also force a greater proportion of individuals into human-wildlife conflict at protected area boundaries (van Eeden *et al*., 2018) as relatively more animals must forage outside of protected areas (Farhadinia *et al*., 2018). It is thus critical that policy actions be well-informed on area requirements of target species. To ensure that protected areas are adequately sized, estimates of the area required for an individual of a given species to persist and reproduce are often quantified via home range analysis (Martins *et al*., 2013, Rechetelo *et al*., 2016, Tédonzong *et al*., 2018). Because occurrence estimators underestimate the area requirements of GPS-tracked animals (often dramatically so), using occurrence estimators to estimate area requirements can result in protected areas that do not accomplish their intended purpose.

Conflating range and occurrence estimators to quantify space use is also dangerous in its implications for basic inference in ecology. For example, the distinction between range and occurrence distributions is particularly salient for studies of resource use and selection by animals. Resource selection is generally studied using resource selection functions—which compare environmental covariates at the locations where animals were present (i.e., “used” locations) to covariates at locations taken from an area assumed to be available for selection (i.e., “available” locations; Manly *et al*. 2007)—or resource utilization functions, which compare intensity of use among an animal’s used locations (Marzluff *et al*., 2004, Millspaugh *et al*., 2006). Range distributions are an appropriate tool for quantifying *availability* for resource *selection* functions, because they characterize the area an animal is likely to travel over the long term. In contrast, occurrence distributions are appropriate for quantifying resource *use* in resource *utilization* functions, because they characterize an animal’s likely presence on the landscape during a study period. In practice, ecologists typically (and correctly) use range estimators to sample availability in resource selection functions, but often use range estimators rather than occurrence estimators to quantify habitat use in resource utilization functions (e.g., Berry *et al*., 2019, Johnston *et al*., 2020, Koizumi & Derocher, 2019, Prince *et al*., 2016, Winder *et al*., 2017). This may be because the initial papers on resource utilization functions (Marzluff *et al*., 2004, Millspaugh *et al*., 2006) used range estimators to generate utilization distributions (understandable because range estimators were the only tools available at the time—occurrence estimators had not been popularized yet). Nevertheless, an increasing number of occurrence estimators have become available over the past two decades (Fleming *et al*., 2016, Horne *et al*., 2007, Johnson *et al*., 2008), and we encourage ecologists to use these occurrence estimators—rather than range estimators—to quantify resource use in resource utilization functions.

Tracking data can and should be a resource for informing our understanding of animal ecology. Although we are now better positioned than ever to use tracking data to estimate different aspects of space use by animals, capturing maximal value from tracking data requires ecologists to understand and use the most rigorous statistical tools and definitions currently available. In this paper, we have highlighted the distinction between range and occurrence distributions, delineated the conditions under which they will behave similarly and differently, mapped ecological questions and statistical estimators to each distribution, and demonstrated the negative consequences of continuing to conflate these two distributions. Both range and occurrence estimators are readily available today in free and open source software (Calabrese *et al*., 2016, Calenge, 2006, Johnson *et al*., 2008, Nielson *et al*., 2013, Signer *et al*., 2019), and we encourage readers to explore the important distinction between range and occurrence estimators themselves (Appendix S1).

## Acknowledgements

We thank S. Alberts, J. Altmann, P. Antunes, J. Goheen, M. Kauffman, A. Paviolo, and M. Xavier da Silva for contributing data for our empirical analyses. J. Calabrese, C. Fleming, and B. Fagan were supported by NSF IIBR 1915347. M. Noonan was supported by an NSERC Discovery Grant RGPIN-2021-02758. F. Jeltsch, N. Blaum and W. Ullmann were supported by the German Research Foundation (DFG) in the framework of the BioMove Research Training Group (DFG-GRK 2118/1). This work was partly funded by the Center for Advanced Systems Understanding (CASUS) which is financed by Germany’s Federal Ministry of Education and Research (BMBF) and by the Saxon Ministry for Science, Culture and Tourism (SMWK) with tax funds on the basis of the budget approved by the Saxon State Parliament.

## Author Contributions

JC conceived the idea; JA, JC, CF, MN, and IS conducted the analyses; and JA and JC led the writing of the manuscript. All authors contributed critically to drafts of the manuscript and gave final approval for publication.

