## Appendix 1 for "Clarifying space use concepts in ecology: range vs. occurrence distributions"

---
title: "Appendix 1: An interactive script demonstrating differences between range and occurrence distribuions"
date: "`r Sys.Date()`"
output: rmarkdown::html_vignette
---


This appendix allows readers to examine differences between range and occurrence distributions by manipulating the characteristics of the data. We begin by fitting a continuous-time movement model on the first 500 locations from a GPS data set from a Cape buffalo (Syncerus caffer) contained in the 'ctmm' R package. Users can use a different data set if they wish--either another data set from 'ctmm' or their own data.
```{r}
# PACKAGES ####
library(ctmm)

# DATA ####
# Using ctmm default data
data(buffalo)
Cilla <- buffalo[[1]]
Cilla <- Cilla[1:500,]
plot(Cilla)

# ANALYSIS ####
# Check out variogram
SVF <- variogram(Cilla)
plot(SVF)

# Guess model fit
GUESS <- ctmm.guess(Cilla, variogram=SVF, interactive=FALSE)

# Fit and select best model
FIT <- ctmm.select(Cilla, GUESS) # takes a few minutes
summary(FIT)
```

The animal's movement track is best characterized by an anisotropic Ornstein-Uhlenbeck Foraging model.

We then create an occurrence distribution that best fits the data conditional on the selected movement model and compare it to the initial data.

```{r}
# Creat an occurrence estimate
OD <- occurrence(Cilla, FIT)

# Compare to the raw data
EXT <- extent(list(Cilla, OD))
par(mfrow=c(1,2))
plot(Cilla, ext=EXT, col.grid=NA, col.level=NA)
title("Location data")
plot(OD, ext=EXT, col.level=NA)
title("Occurrence UD")
```

The areas of high probability adhere closely to the movement track.

We then create a range estimate that best fits the data conditional on the selected movement model and compare it to the occurence estimate.

```{r}
# Create a range estimate
RD <- akde(Cilla, FIT)

# Comparison plot with AKDE
EXT <- extent(list(RD,OD))
par(mfrow=1:2)
plot(RD, ext=EXT, col.grid=NA, col.level=NA)
title("Range UD")
plot(OD, ext=EXT, col.level=NA)
title("Occurrence UD")
```

When compared to the occurrence estimate, the range estimate adheres much less tightly to the movement track.

We then manipulate the data so that every other location is deleted and make both a range and occurrence estimate. We then compare the new estimates to the initial estimates created using the full movement track.

```{r}
# Manipulation 1 - Coarsening the data
SUB <- buffalo$Cilla

SUB <- Cilla[rep(c(TRUE,FALSE),length.out=nrow(Cilla)),]
RD.SUB <- akde(SUB, FIT)
OD.SUB <- occurrence(SUB, FIT)
par(mfrow=c(2,2))
plot(RD, ext=EXT, col.grid=NA, col.level=NA)
title("Range UD")
plot(OD, ext=EXT, col.level=NA)
title("Occurrence UD")
plot(RD.SUB, ext=EXT, col.grid=NA, col.level=NA)
title("Range UD (Subset)")
plot(OD.SUB, ext=EXT, col.level=NA)
title("Occurrence UD (Subset)")
```

You can see that while the new range estimate on the coarsened data looks nearly identical to the original range estimate, the new occurrence estimate has a much larger area than the original occurrence estimate.

We then manipulate the data again so that the data set is half as long and make both a range and occurrence estimate. We then compare this second set of new estimates to the initial estimates created using the full movement track.

```{r}
# Manipulation 2: Truncating the data
SUB2 <- Cilla[1:(nrow(Cilla)/2),]

RD.SUB2 <- akde(SUB2, FIT)
OD.SUB2 <- occurrence(SUB2, FIT)

# Create graphs
par(mfrow=c(2,2))
plot(RD, ext=EXT, col.grid=NA, col.level=NA)
title("Range UD")
plot(OD, ext=EXT, col.level=NA)
title("Occurrence UD")
plot(RD.SUB2,ext=EXT,col.grid=NA,col.level=NA)
title("Range UD (Subset 2)")
plot(OD.SUB2,ext=EXT,col.level=NA)
title("Occurrence UD (Subset 2)")
```

You can see that while the new range estimate on the truncated data again looks quite similar to the original range estimate, the new occurrence estimate now has a much *smaller* area than the original occurrence estimate.

These simple examples demonstrate that (1) range and occurrence estimates can be very different from one another, and (2) occurrence estimates can change substantially with relatively small differences in data, while range estimates remain much more stable. Home range estimates generated using occurrence estimators will often be both too small and much more susceptible to sampling scheme of the underlying data set.
